# scMoC: Single-Cell Multi-omics clustering

**DOI:** 10.1101/2021.02.24.432644

**Authors:** Mostafa Eltager, Tamim Abdelaal, Ahmed Mahfouz, Marcel J.T. Reinders

## Abstract

**Motivation:** Single-cell multi-omics assays simultaneously measure different molecular features from the same cell. A key question is how to benefit from the complementary data available and perform cross-modal clustering of cells.

**Results:** We propose **S**ingle-**C**ell **M**ulti-**o**mics **C**lustering (scMoC), an approach to identify cell clusters from data with co-measurements of scRNA-seq and scATAC-seq from the same cell. We overcome the high sparsity of the scATAC-seq data by using an imputation strategy that exploits the less-sparse scRNA-seq data available from the same cell. Subsequently, scMoC identifies clusters of cells by merging clusterings derived from both data domains individually. We tested scMoC on datasets generated using different protocols with variable data sparsity levels. We show that, due to its imputation scheme, scMoC 1) is able to generate informative scATAC-seq data due to its RNA guided imputation strategy, and 2) results in integrated clusters based on both RNA and ATAC information that are biologically meaningful either from the RNA or from the ATAC perspective.

**Availability:** The code is freely available at: https://github.com/meltager/scmoc.

**Supplementary information:** Supplementary data are available online.

## 1 Introduction

Recent development in single-cell technologies have enabled measuring different molecular features with increasing throughput. However, due to the destructive nature of most protocols, these molecular features can be measured from the same biological sample but from different cells. Recent advances have made it possible to profile more than one ‘omic’ data from the same cell, introducing single-cell multimodal omics (Zhu *et al*. 2020). Such technology enables simultaneous measurement of, for example, gene expression (scRNA-seq) and chromatin accessibility (scATAC-seq), as in the sci-CAR protocol (Cao *et al*. 2018). It is anticipated that single-cell multimodal omics is a promising technology that will improve our ability to dissect the complex gene regulatory networks, cell lineages and trajectories)Schier 2020).

Several methods are proposed to integrate different omics data but from unpaired datasets, i.e., measured from different cells but from the same biological sample. For example, Seurat V3 (Stuart *et al*. 2019) and LIGER (Welch *et al*. 2019) map the common feature space into an aligned latent domain using dimensionality reduction. Nevertheless, single-cell multimodal omics brings up a different computational challenge by offering *paired data* points for the same cell (Anjun *et al* 2020; Stuart and Satija 2019; Zhu *et al* 2020).

One of the main challenges lies in the data sparsity. For example, the data density in (unimodal) scRNA-seq datasets usually ranges from 10% to 45% and from 1% to 10% in (unimodal) scATAC-seq datasets (Chen *et al* 2019). But, when doing multimodal omics measurements these densities lower considerably to 1% -10% for the scRNA-seq data and to 0.2 % – 6% for scATAC-seq. This can be accounted for to the prematurity of the protocols and modalities (Stuart and Satija 2019). Importantly, data sparsity can deteriorate cluster separation and, thus, make it hard to capture a clear structure in the cellular composition (Baek and Lee 2020; Luecken *et al* 2020).

The cluster agreement between data from different modalities represents another challenge. Despite that measurements of different modalities are taken from the same cells, they relate to different functionalities within a cell. Hence, when approached individually, these measurements will not result in the same grouping of cells, complicating the decision on a correct grouping as well as the establishment of cell types. So, this calls for algorithms that can exploit the complementary view on the same cell (Stuart and Satija 2019).

To tackle these challenges, we propose **S**ingle-**C**ell **M**ulti-**o**mics **C**lustering (scMoC). scMoC is designed to cluster paired multimodal datasets that measures both single-cell transcriptomics sequencing (scRNA-seq) and single-cell transposase accessibility chromatin sequencing (scATAC-seq). The most important ingredient of scMoC is that it imputes the sparse scATA-seq data using an RNA-guided imputation process. A *k-nn* based imputation imputes ATAC peak counts for a specific cell from a set of similar cells by, for example, the average of the peak counts found in the set of similar cells. This imputation scheme relies on the set of similar cells, which are hard to find when data sparsity is high as that impacts the resemblance of peak profiles. By recognizing that we have measured RNA counts from the same cells and that the RNA data sparsity is much lower, we propose to calculate cell similarities not in the ATAC domain, but in the RNA domain. We show that the resulting RNA-guided imputed ATAC data is better structured and provides informative and complementary data in comparison to analysis based on the RNA data only.

## 2 Methods

### 2.1 scMoC

A general overview of scMoC is shown in Figure 1. Briefly, to overcome the data sparsity in the scATAC-seq data, we impute the scATAC-seq data. Here, we use the fact that the data is paired and guide the scATAC-seq imputation by choosing similar cells based on the RNA measurements of the cell. Next, we cluster the scRNA-seq and the imputed scATAC-seq independently and merge the resulting clusters. In the following sections, we describe the details of the method. A more precise workflow can be found in Supplementary Figure 1.

**Figure 1:**
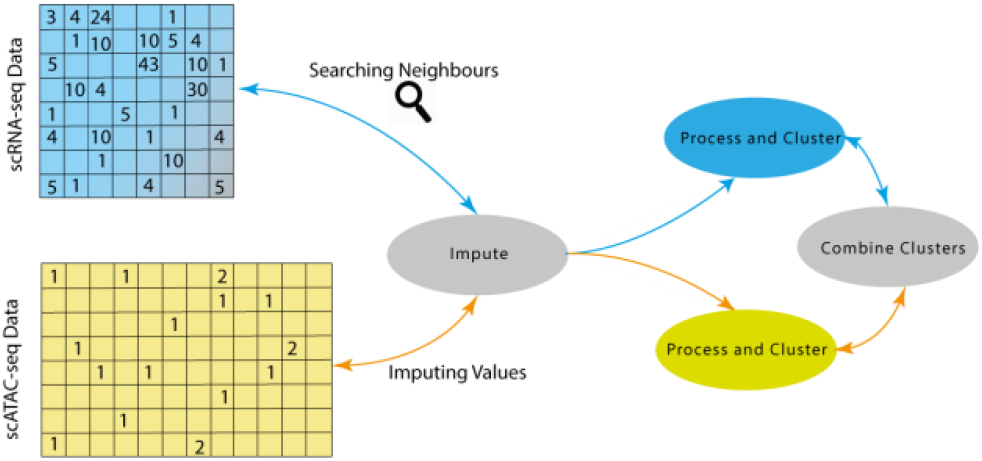
Schematic overview of scMoC. scMoC clusters multi-modal single-cell data based on scRNA-seq and scATAC-seq measurements from the same cell. It encompasses an RNA-guided imputation strategy to leverage the higher data sparsity of the scRNA-seq data (with respect to the scATAC-seq data). scMOC builds on the idea that cell-cell similarities can be better estimated from the RNA profiles and then used to define a neighborhood to impute from it the ATAC data, since these are co-measured from the same cell. After the imputation, the two modalities are clustered individually and then combined into one clustering in which RNA-based clusters are being split if there is enough evidence from the ATAC data.

#### 2.1.1. Pre-processing of the data

scRNA-seq data is pre-processed following the Seurat 3.0 pipeline (Stuart *et al*. 2019). For the scRNA-seq data, first, a quality control step is performed which removed noisy and almost empty cells. These thresholds are database dependent (Datasets and Supplementary Table 1). scRNA-seq counts are normalized by dividing each count by the total count for each cell and then scaling up using a factor of 1e4, followed by a log(*x* + 1) transformation. The top 6,000 highly variable genes are selected, using the variance stabilizing transform (vst) algorithm. Subsequently, the data is centered around zero and scaled by dividing each expression by the standard deviation. Next, we projected the data on the top 20 Principal Components for further processing.

For the scATAC-seq data, we followed common pipeline steps (Baek and Lee 2020). First, we removed almost empty and noisy cells (Dataset and Supplementary Table 1). Peak counts are normalized by dividing each count by the total count for each cell and then scaling up using a factor of 1e4. This is followed by an imputation step (next section). After imputation, the scATAC-seq data is renormalized and log(*x* + 1) transformed. Then the data is scaled and Latent Semantic Indexing (LSI) is used to reduce the dimensionality to the top 20 LSI vectors for further processing.

#### 2.1.2 Imputation of the scATAC-seq data

To deal with the sparsity of the scATAC-seq data, we resort to an imputation strategy. Here, we exploit the fact that for the same cell we have RNA and ATAC measurements. Hence, we propose to guide the imputation of the scATAC-seq data using the scRNA-seq data because cell-cell similarities can be calculated more robustly on the scRNA-seq data as it suffers less from data sparsity. For this *RNA-guided imputation*, a set of similar cells are chosen for each cell. Hereto, we choose the top-*k* (*k*=50 throughout this work) closest cells based on the Euclidean distance between cells in the PCA projected space of the RNA measurements. scATAC-seq peak values are then imputed by taking the average peak value across the *k*-nearest neighboring cells.

For reference, we also considered a *self-imputation* in which the *k*-nearest neighboring cells are selected on the basis of the scATAC-seq data only, using the Euclidean distance of the unimputed scATAC-seq data projected in the LSI space as a measurement of the distance between cells. The imputation method afterwards remains the same as mentioned in the *RNA-guided imputation*.

#### 2.1.3 Unimodal clustering

Both, the scRNA-seq and (imputed) scATAC-seq data are clustered separately using the Louvain algorithm (Blondel et al. 2008), implemented in Seurat with graph resolution of 0.8. To build the neighborhood graph, we used the *k*-nearest neighbor algorithm (with *k*=50) and using the Euclidean distance.

#### 2.1.4 Combining RNA and ATAC clustering

To combine the scRNA-seq and (imputed) scATAC-seq based clusters, we (again) use a strategy that trust the scRNA-seq rich data more than the scATA-seq data. Hence, we choose only to split RNA-based clusters when there is enough evidence on from the scATAC-seq data (and not the other way around). To do so, we construct a contingency table representing the percentage of agreement between both clustering. When an RNA cluster overlaps with more than one ATAC cluster, scMoC splits the RNA cluster accordingly, i.e. if the overlapping percentage of the ATAC cluster is larger than 10% and less than 90% of the RNA cluster, we accept that split. After splitting, the non-assigned cells from the original RNA cluster are assigned to the closest cluster based on the average distance to the cells within a cluster using Euclidean distance in the RNA space only.

### 2.2 Datasets

Three different datasets were used to test scMoC (summarized in Table 1): *mouse kidney data*, measured using the sci-CAR protocol (Cao *et al*. 2018); *adult mouse cerebral cortex data*, measured using the SNARE-seq protocol (Chen *et al*. 2019); and *human peripheral blood mononuclear cells (PBMC) data* from 10x Multiome. These datasets were chosen because they range in their data density; about 6-fold for the scRNA-seq data and more than 20-fold for the ATAC data. For the quality control step, each dataset is filtered differently based on the quality control metric visualization tools provided in Seurat package (Stuart *et al*. 2019). Supplementary Table 1 list the limits used in this step.

**Table 1.** Description of the datasets used in this study.

|  | sci-CAR | SNARE-seq | 10X genomics |
| --- | --- | --- | --- |
| Tissue | Mouse Kidney | Mouse brain | PBMC |
| # of Cells | 11,296 | 10,309 | 3,003 |
| # of Genes (RNA) | 49,584 | 33,160 | 21,547 |
| # of Peaks (ATAC) | 252,741 | 244,544 | 75,540 |
| RNA data density | 1.05% | 2.87% | 7.06% |
| ATAC data density | 0.28% | 1.01% | 6.49% |

## 3 Results

### 3.1 scMoC reveals new cell clusters from the data

#### RNA-guided imputation improves ATAC data quality

We tested scMoC on the sci-CAR dataset. Figure 2A shows the UMAP for the scRNA-seq part of the data only as well as the identified RNA-based clusters (Methods), which shows that the clusters of cells are clearly separable. This is not the case for the scATAC-seq part of the data (Figure 2B), also resulting in low cluster agreement between the two clusterings (see contingency matrix, Supplementary Figure 2). Probably that is driven by the high sparsity in the scATAC-seq data (here 99.7% zeros, see Table 1), largely influencing distances between cells in the ATAC data. This can be resolved by imputing the ATAC data. Hereto, we applied our proposed RNA-guided imputation (Methods), which decreased the data sparsity to 89% zeros. The resulting imputed ATAC data indeed shows well-separated clusters (Figure 2C), as well as improved cluster agreements (Supplementary Figure 3). For comparison, we imputed the ATAC data with the self-imputed scheme (using only the ATAC data, Methods), which slightly reduced the data sparsity (towards 95.9%) and did not result in separable clusters (Figure 2D), as well as a low cluster agreement with the RNA-only based clusters (Supplementary Figure 4).

**Figure 2:**
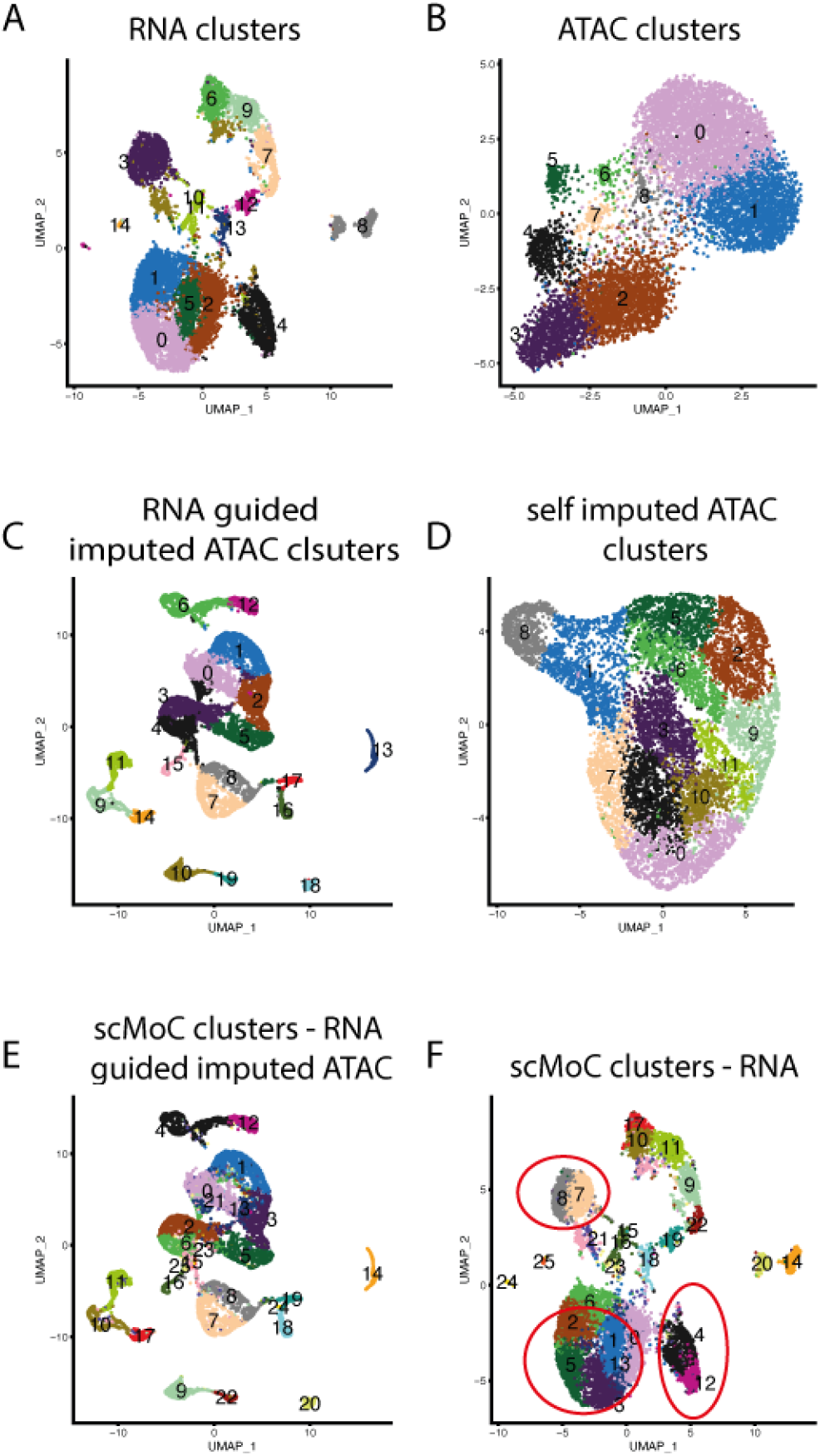
scMoC shows different cluster in the scRNA-seq data based on the imputed scATAC-seq data. UMAP visualizations of the sci-CAR. (A) scRNA-seq data, colors indicating RNA-based clusters. (B) scATAC-seq data, colors indicating ATAC-based clusters. (C) RNA-guided imputed scATAC-seq data, colors indicating clusters within the imputed data. (D) Self-imputed scATAC-seq data, colors indicating clusters detected in this data. (E) RNA-guided imputed scATAC-seq data colored according to the scMoC clusters. (F) scRNA-seq data colored according to the scMoC clusters.

#### ATAC-based clusters refine RNA-based cluster

Next, we combined the RNA-guided imputed ATAC-based clusters with the RNA-based clusters using scMoC (Methods). Figure 2E and Figure 2F show the resulting scMoC clusters overlaid on the UMAPs of the scRNA-seq and RNA-guided imputed scATAC-seq data, respectively. This shows that the original RNA-based clusters are split based on the scATAC-seq data. For example, the red circles in Figure 2F indicate splits of the RNA-based clustering due to the available ATAC data of these same cells.

#### ATAC refined clusters provide complementary information

To ensure that scMoC clusters cannot be achieved by just tweaking the clustering resolution, we compared the scMoC clusters to more fine-grained clusterings of, both, the scRNA-seq and the imputed scATAC-seq data. Figure 3 shows how the different clusterings are related. For example, scMoC cluster 22 (scMoC_22) overlaps with scRNA-seq cluster 27 (RNA_27) as well as the scATAC-seq based cluster 31 (ATAC_13), indicating an agreement of the grouping of cells when looking only at RNA or ATAC based information. Alternatively, scMoC cluster 5 (scMoC_5) overlaps with scRNA-seq cluster 10 (RNA_10) and combines two scATAC-seq based clusters (ATAC_16, ATAC_18). This nicely shows that the ATAC data provides additional information to split the RNA cluster. Interesting more complex combinations do occur. For example, scMoC cluster 0 (scMoC_0) combines two scRNA-seq clusters (RNA_2 and RNA_7) and (parts of) four scATAC-seq based clusters (ATAC_4, ATAC_5, ATAC_19 and ATAC_23). This complex combination is further supported the contingency table between the scRNA-seq and RNA-guided imputed scATAC-seq clusters as there is no one-to-one agreement between these clusters (see contingency matrix, Supplementary Figure 5).

**Figure 3:**
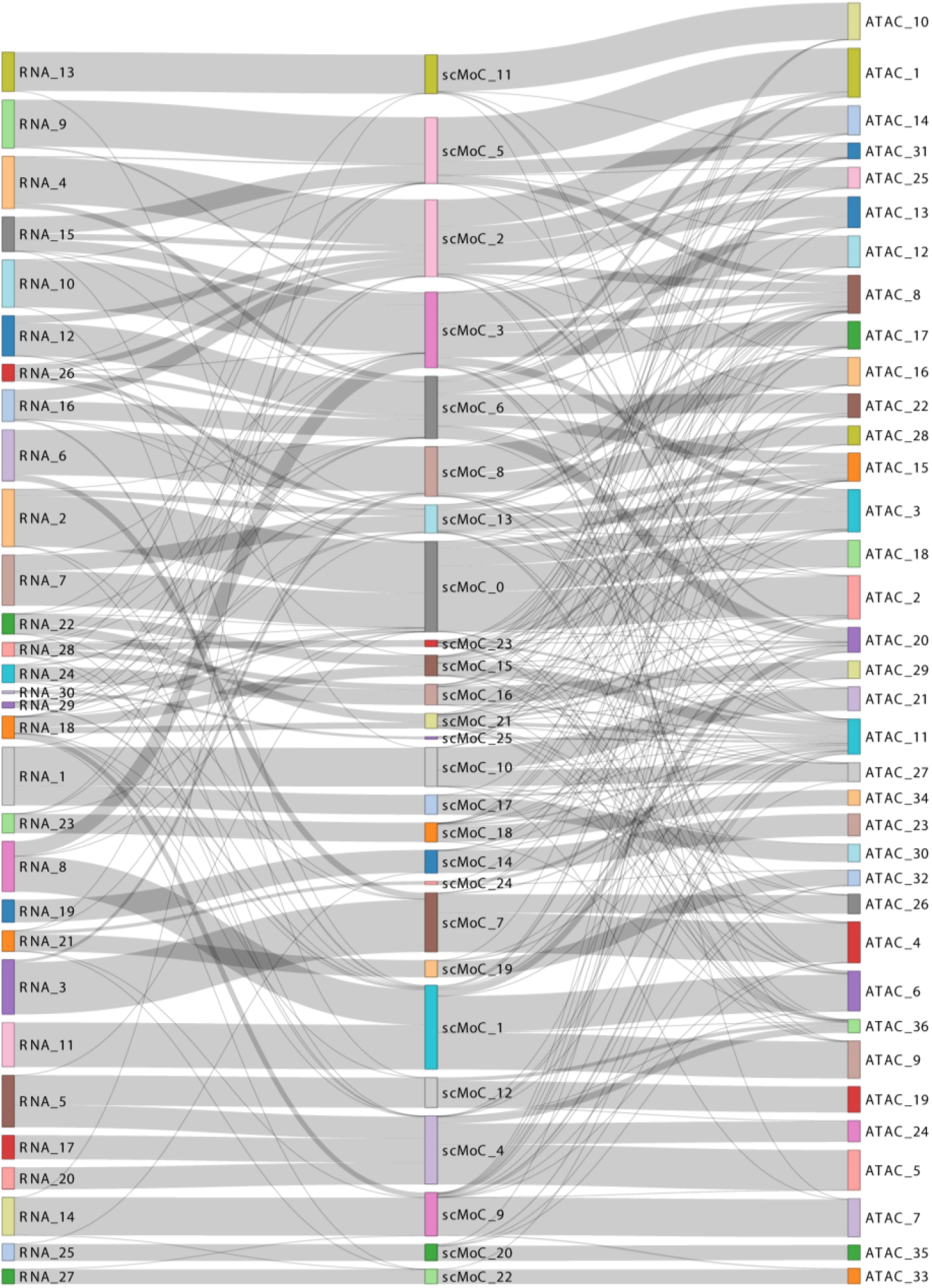
Sankey graph showing how the scMoC clustering relates to the scRNA-seq and (imputed) scATAC-seq based clusterings. The middle panel shows the detected scMoC clusters. To show that the ATAC-based splits induced (of the scRNA-seq clustering) are not similar to a more fine-grained clustering of the RNA data. The left panel shows the RNA-based clusters when using a more detailed resolution (so more RNA clusters). Same holds for the right panel showing the fine-grained clusters of the ATAC data. In some cases (e.g. the top scMoC cluster 11), the more fine-grained RNA clustering did not split the RNA cluster, as can be noticed that scMoC cluster 11 completely overlaps with the more fine-grained RNA cluster 12. For scMoC cluster 1 (scMoC_1), we see that it is split into clustters RNA_19 and RNA 21 when we cluster at a more fine-grained level. We also observe that this splitting matches a more fine-grained clustering of the ATAC data, that is RNA cluster RNA_19 matches the more fine-grained ATAC cluster ATAC_6 and, similarly, RNA cluster RNA_11 matches ATAC cluster ATAC_9. Such agreements do, however, not always occur. For example, scMoC cluster C splits partly in RNA clusters RNA_8 which resembles ATAC cluster 6, whereas on the higher level it resembles ATAC cluster 13, indicating that indeed the scATAC-seq and scRNA-seq data give complementary information

### 3.2 Biological interpretation for scMoC clusters

#### Genes differentiating scMoC clusters overlap with cell type marker genes

To explore the biological relevance of the resulting scMoC clusters, we calculated the differentially expressed (DE) genes (Wilcoxon rank-sum test and Bonferroni for multiple test correction) for each cluster (against all other clusters) and checked if the list of DE genes includes well-known marker genes for mouse kidney cell types (http://bio-bigdata.hrbmu.edu.cn/CellMarker/). Figure 4 depicts a dot plot indicating the association of the marker genes with each of the clusters. For example, *Cndp2, Cyp2e1, Keg1*, S*lc13a3, Ugt2b38* are differentially expressed in cluster 1 and are marker genes for Proximal tubule cells. Cluster 10 represents Distal convoluted tubule cells as its DE genes overlap with the markers: *Abca13, Calb1, Sgms2, Slc12a3, Slc16a7, Trpm6, Trpm7, Tsc22d2*, and *Wnk1*. While cluster 14 is associated with Collecting duct intercalated cell marker genes (e.g. *Atp6v0d2, Atp6v1c2, Car12, Insrr, Ralgapa2, Rcan2, Slc26a4, Slc4a9, Syn2, Tmem117*). A full list for marker genes and clusters can be found in Supplementary Figure 6.

**Figure 4:**
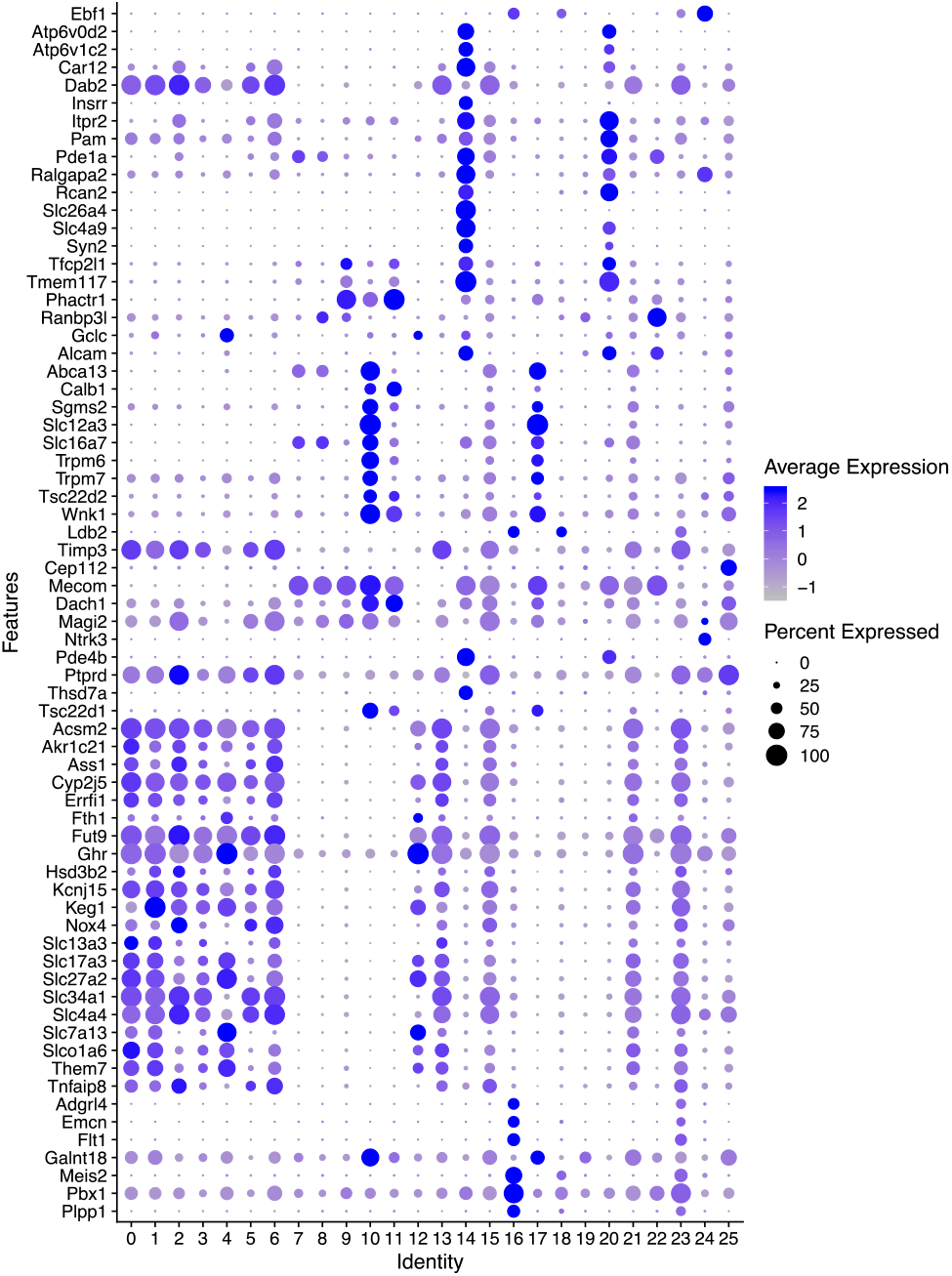
scMoC clusters marker genes. Dot plot showing the differentially expressed genes (i.e. markers) of the scMoC clusters. Expression of a set of Mouse kidney marker genes across scMoC clusters. The color intensity of the dot represents the average expression of the gene across the cells in the cluster and the size relates to the percentage of cells within the cluster expressing each gene.

#### ATAC induced splits do show differences in marker genes

The dot plot shows that some clusters have almost the same list of marker genes. For example, clusters 15 and 19. These two clusters originate from one scRNA-seq based cluster, but are split based on the scATAC-seq data. Interestingly, we do observe that there are differences in the level of the expressions of the associated marker genes between the two scMoC clusters. However, these differences were not sufficient to split in two clusters based on the RNA data alone. But, incorporating the imputed ATAC data boosts the power to differentiate between these two subpopulations.

#### Differentially expressed genes for ATAC induced splits

To study induced cluster splits more thoroughly, we calculated the differential expression between splitted clusters. Figure 5 shows the volcano plot of such a split between scMoC clusters 3 and 5, which originally were one RNA-only cluster (Cluster 0, see Figure 2A). Both clusters are categorized as Proximal tubule cell as derived from the associated marker genes (Figure 4). But, Figure 5 shows that the two scMoC clusters have a different distribution for the identified marker genes. Supplementary Figure 7 shows similar observations for other cluster splits.

**Figure 5:**
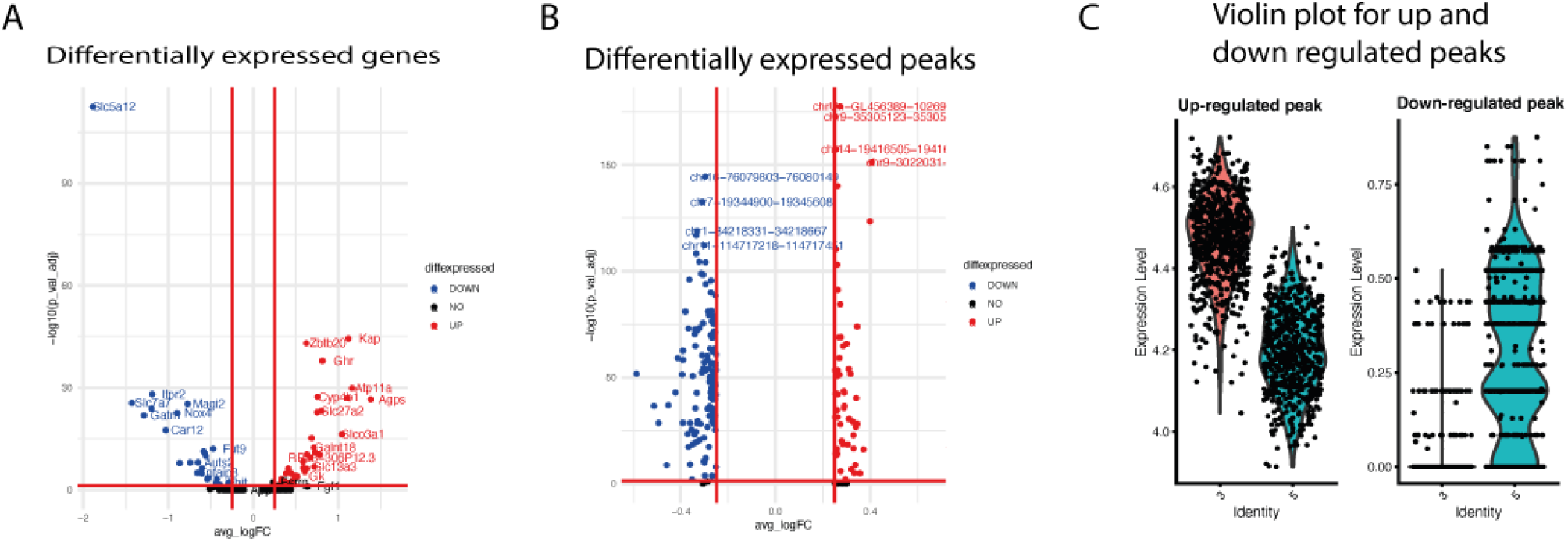
Differentially expressed genes and peaks for ATAC induced splits. Volcano plots showng (A) differentially expressed genes, and (B) between differentially express ATAC peaks between scMoC cluster 3 and scMoC cluster 5. (C) Violin plot for the top up-regulated peak (i.e. chrUn-GL456389-10269-10904) and top down-regulated peak (i.e. chr16-76079803-76080149).

#### ATAC peaks mark ATAC induced splits

To check the effect of scMoC clusters on the ATAC data, and to find whether the scMoC clusters preserve the cluster differences. We calculated the differential expressed peaks between scMoC clusters 3 and 5. Figure 5B shows the difference in the expression of the peaks between these two clusters. We selected the top up-regulated and down-regulated peaks between these clusters and showed the difference in the distribution between these clusters cells (Figure 4C).

### 3.3 Applying scMoC to different protocols

#### scMoC generalizes to other protocols suffering from data sparsity

To test how well the proposed scMoC pipeline deals with another multimodal protocol suffering from data sparsity, we applied it to SNARE-seq data that has a data sparsity of 99% approximately. The RNA guided imputation decreased the data sparsity to 77.6% (Figure 6B and 6D). Combining the imputed ATAC-only clustering with the RNA-only (Figure 6A) clustering lead to new clusters (Figure 6C and 6E) which were not observed either in the RNA-only or the unimputed ATAC-only analysis.

**Figure 6:**
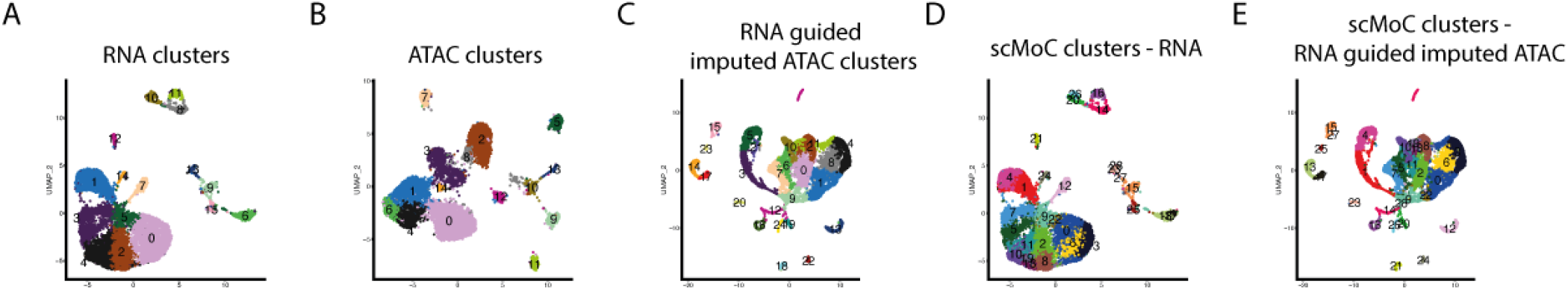
Applying scMoC to SNARE-seq data. UMAP visulation for SNARE-seq data. (A) scRNA-seq data, colors indicating the RNA-based clusters. (B) scATAC-seq data, colors indicating ATAC-based clusters. (C) RNA-guided imputed scATAC-seq data, colors indicating clusters within the imputed data clustred independanlty. (D) scRNA-seq data colored according to the scMoC clusters. (E) RNA-guided imputed scATAC-seq data colored according to the scMoC clusters

#### scMoC does not apply when data sparsity becomes similar to that of the scRNA-seq

Next, we applied scMoC to 10X genomics multiome data for which the scATAC-seq data has about the same sparsity as the scRNA-seq data (both about 93%). Figure 7A and 7B display the scRNA-seq and scATAC-seq data, showing that the scATAC-seq data has a similar clustering structure as that of the scRNA-seq data (Supplementary Figure 8). When imputing the ATAC data using scMoC, Figure 7C, the distribution of the cells is clearly affected, potentially confusing the grouping of cells based on the (imputed) ATAC data.

**Figure 7:**
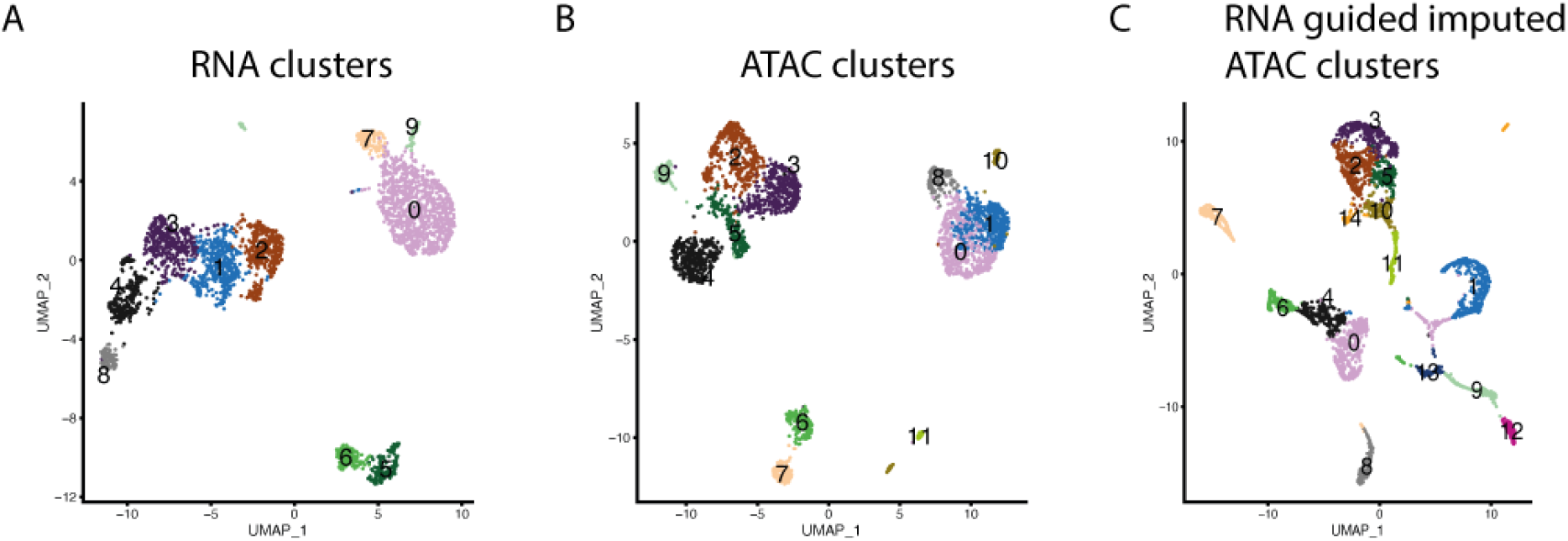
Applying scMoC to 10X genomics multiome data. UMAP visualization (A) and (B) the RNA and the ATAC, respectively. (C) RNA guided imputed ATAC data.

## 4 Discussion

We have shown the ability for scMoC to co-cluster data from scRNA-seq and scATAC-seq measured from the same cell. scMoC’s imputation strategy made it possible to exploit the data sparse scATAC-seq data. scMoC’s joint clustering scheme of both data domains revealed biologically meaningful clusters that are supported by the expression of known cell-type-specific markers. An important contribution of scMoC is its imputation strategy: it uses the less data sparse modality (here, the scRNA-seq data) to impute the data sparse modality (here, the scATAC-seq data). This leverages the fact that both modalities are measured at the same cell; and thus that the other data modality can thus be used to find similar cells. We have shown that when the differences in data sparsity between the two data modalities decreases (such as in the 10X-genomics data), the need for such an imputation scheme also decreases. With our work we have highlighted that single cell multimodal measurements are a valuable tool to resolve heterogeneity at the single cell level, even if the data sparsity of one of the modalities is low.

## Funding

European Union’s H2020 research and innovation programme under the MSCA grant agreement [861190 (PAVE)]; European Commission of a H2020 MSCA award [675743] (ISPIC); NWO Gravitation project: BRAINSCAPES: A Roadmap from Neurogenetics to Neurobiology (NWO: 024.004.012).

## Conflict of interest statement

None declared.

REFERENCE

## Conflict of Interest

none declared.

## Notes

### Competing Interest Statement

The authors have declared no competing interest.

