## Supplementary figures and tables for "scMoC: Single-Cell Multi-omics clustering"

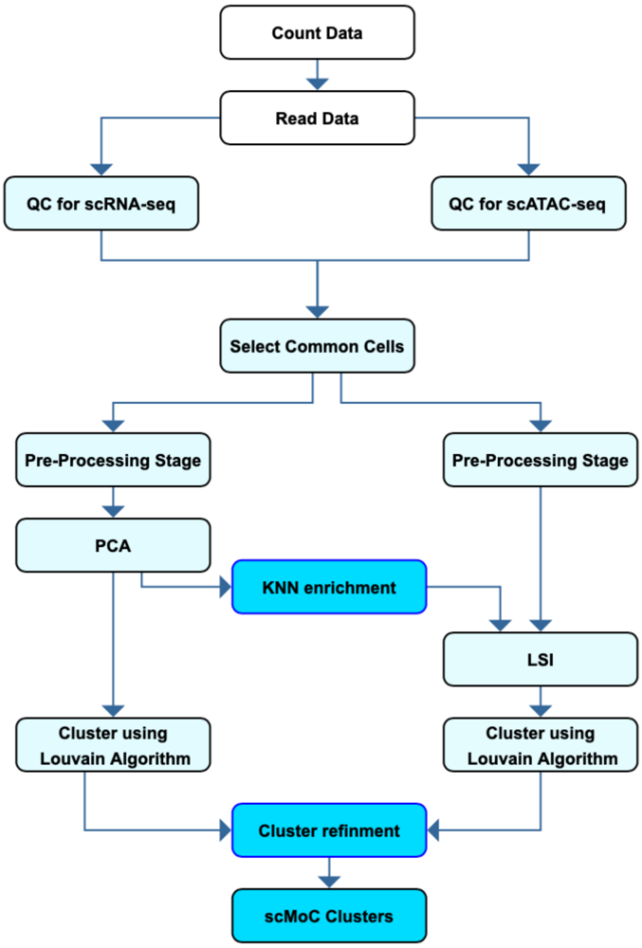


**Supplementary Figure 1: scMoC block diagram** scMoC applies the Seurat V3 pipeline in processing the RNA data. First The Data undergoes a quality control step. Then the cells that have both RNA and ATAC data measured and passed the previous step is selected to go through the rest of the steps. For the RNA, the data undergoes the pre-processing stage of Normalizing the data and then scaling up by factor 1e4. The Data is then projected to the PCA space. Afterwards, the data RNA data is clustered. For the ATAC data after the pre-processing stage, the neighborhood of each cell is searched in the PCA projected space of the RNA. The KNN enrichment step is to take the average of the peak values for each cell. After that the data is renormalized and projected to the LSI space to be clustered. After clustering both data domains the cluster refinement step is made to get the splits of the ATAC in the RNA cluster and then he non-assigned cells from the original RNA cluster are assigned to the closest cluster based on the average distance to the cells within a clus-ter using Euclidean distance in the RNA space only.

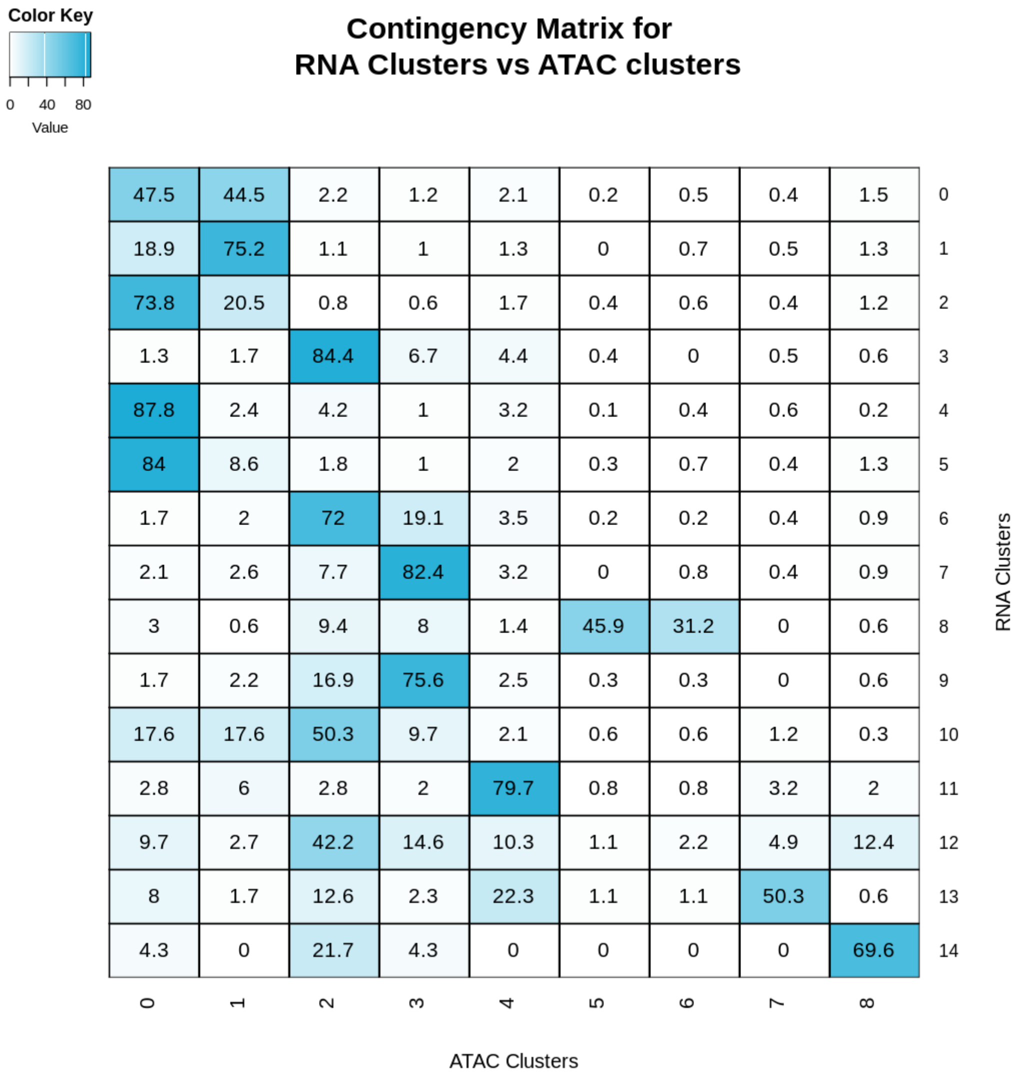


**Supplementary Figure 2 :** The Contingency Matrix between the scRNA-seq and scATAC-seq without imputation,

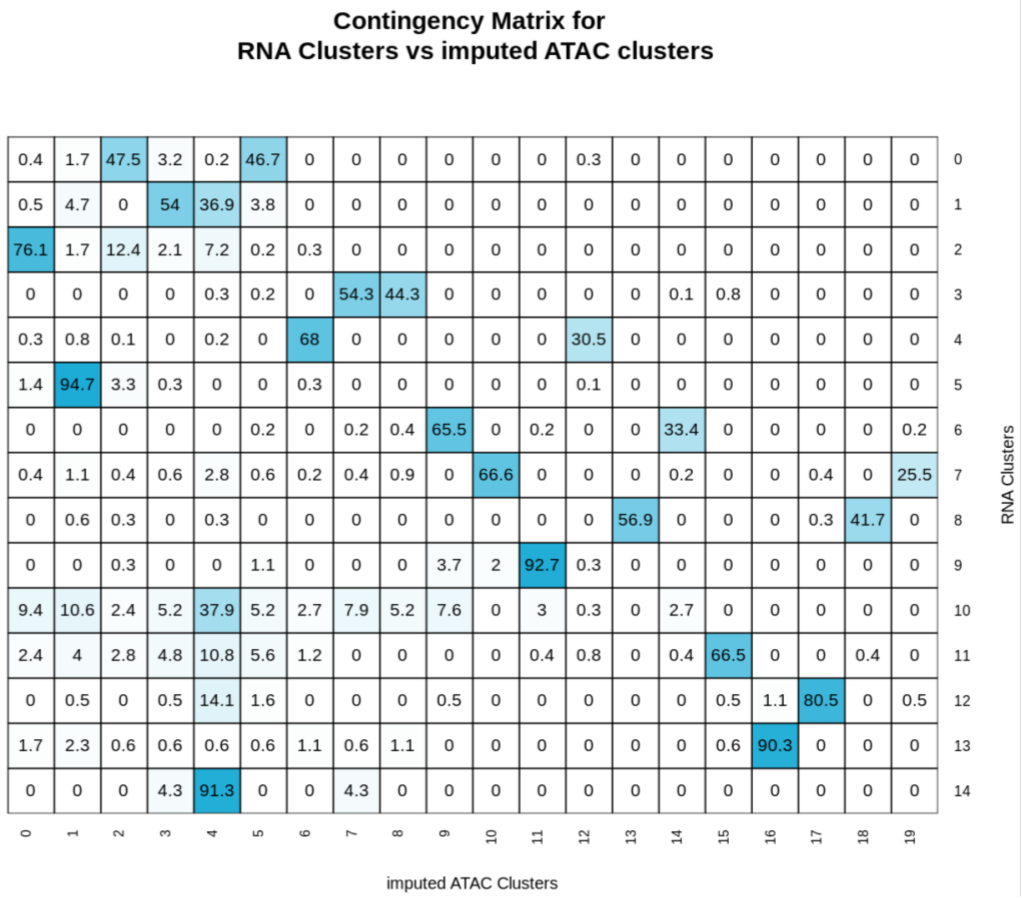


**Supplementary Figure 2** shows the Contingency Matrix between the scRNA-seq and scATAC-seq with RNA guided imputation. This matrix is used to decide on the splitting and refinement of the scMoC clustering. When there is an agreement between the clusters more thant 10% and

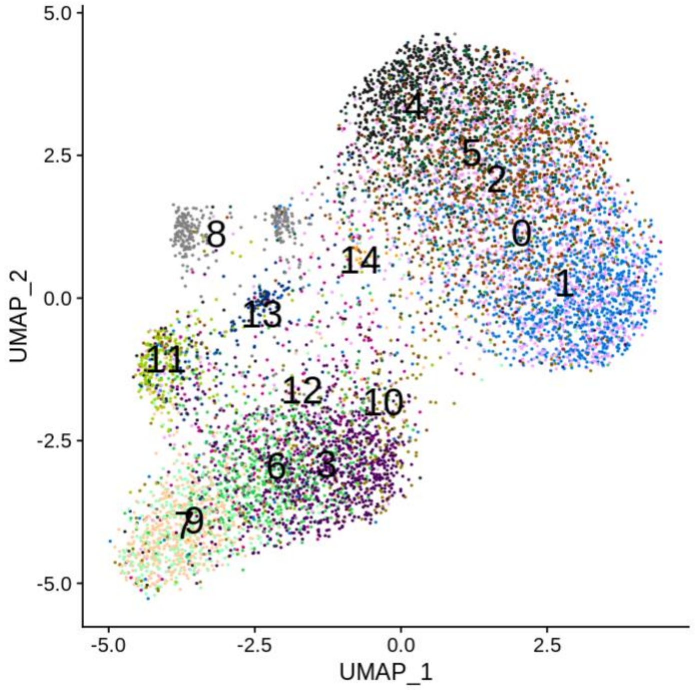


**Supplementary Figure 4:** sci-CAR scATAC-seq data colored with the scRNA-seq clusters showing a discrepancy between both clustering in both data domains

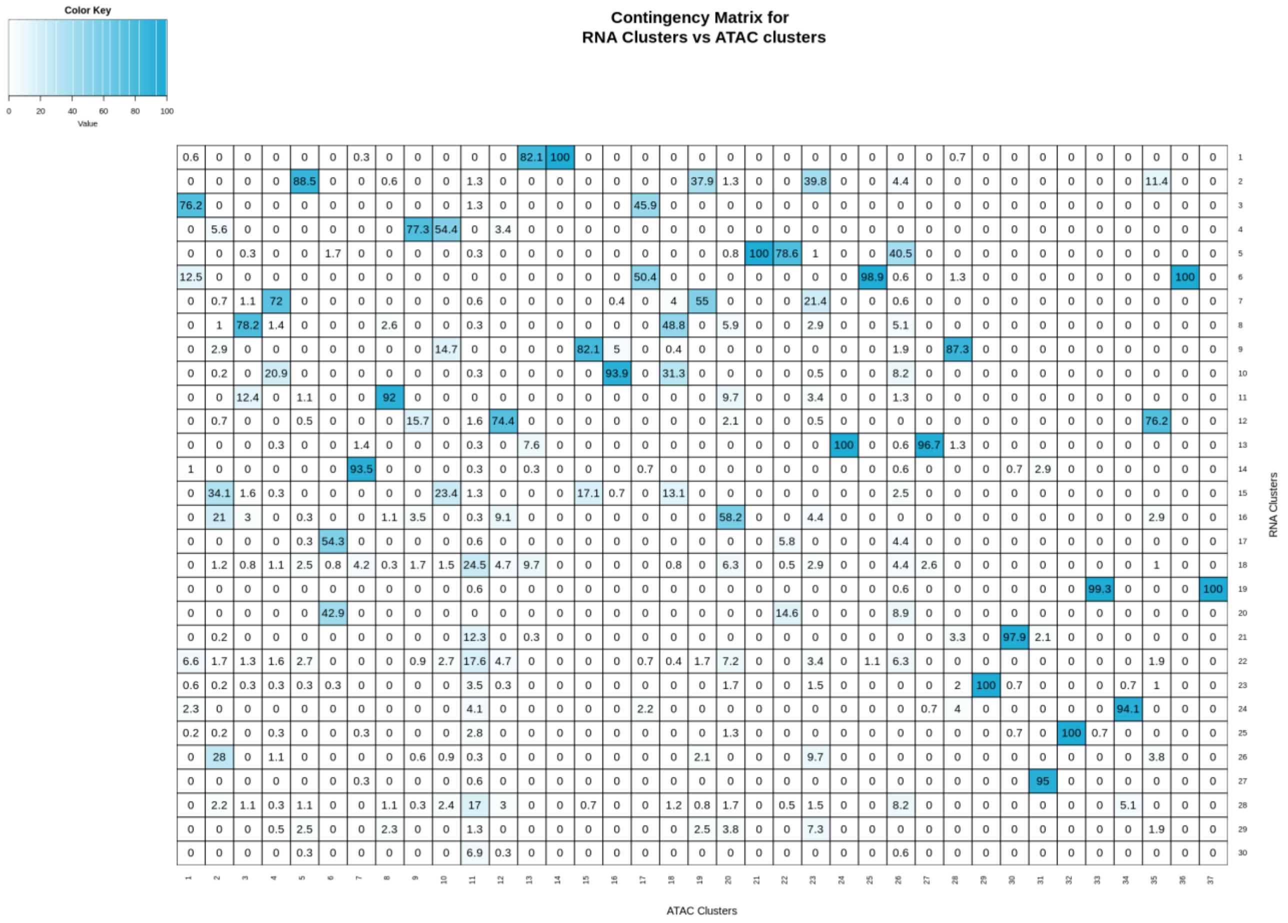


**Supplementary Figure 5 :** Contingency table for the sci-CAR data between scRNA-seq and scATAC-seq data clustered with higher level of resolution in the clustering algorithm to generate double the number of clusters in the scRNA-seq data. This shows that there is no one-to-one agreement between he clusters.

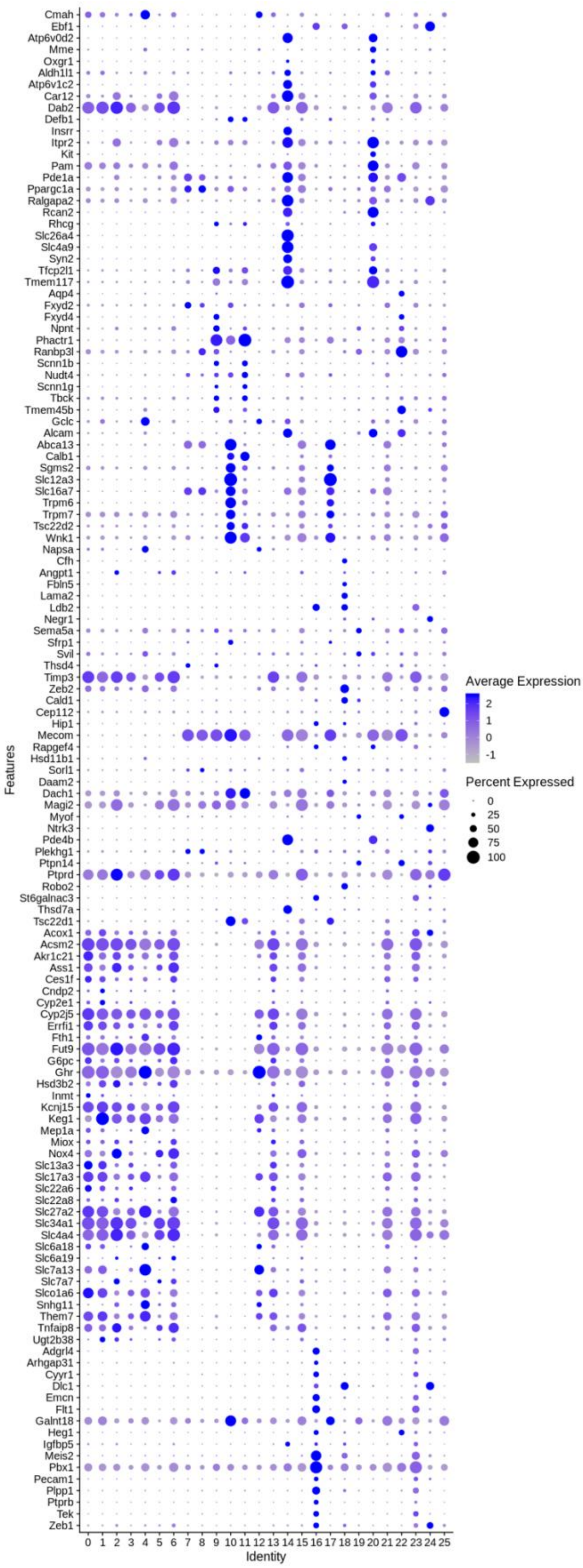


**Supplementary Figure 6 :** Full list of the marker genes that are found in the scMoC clusters.

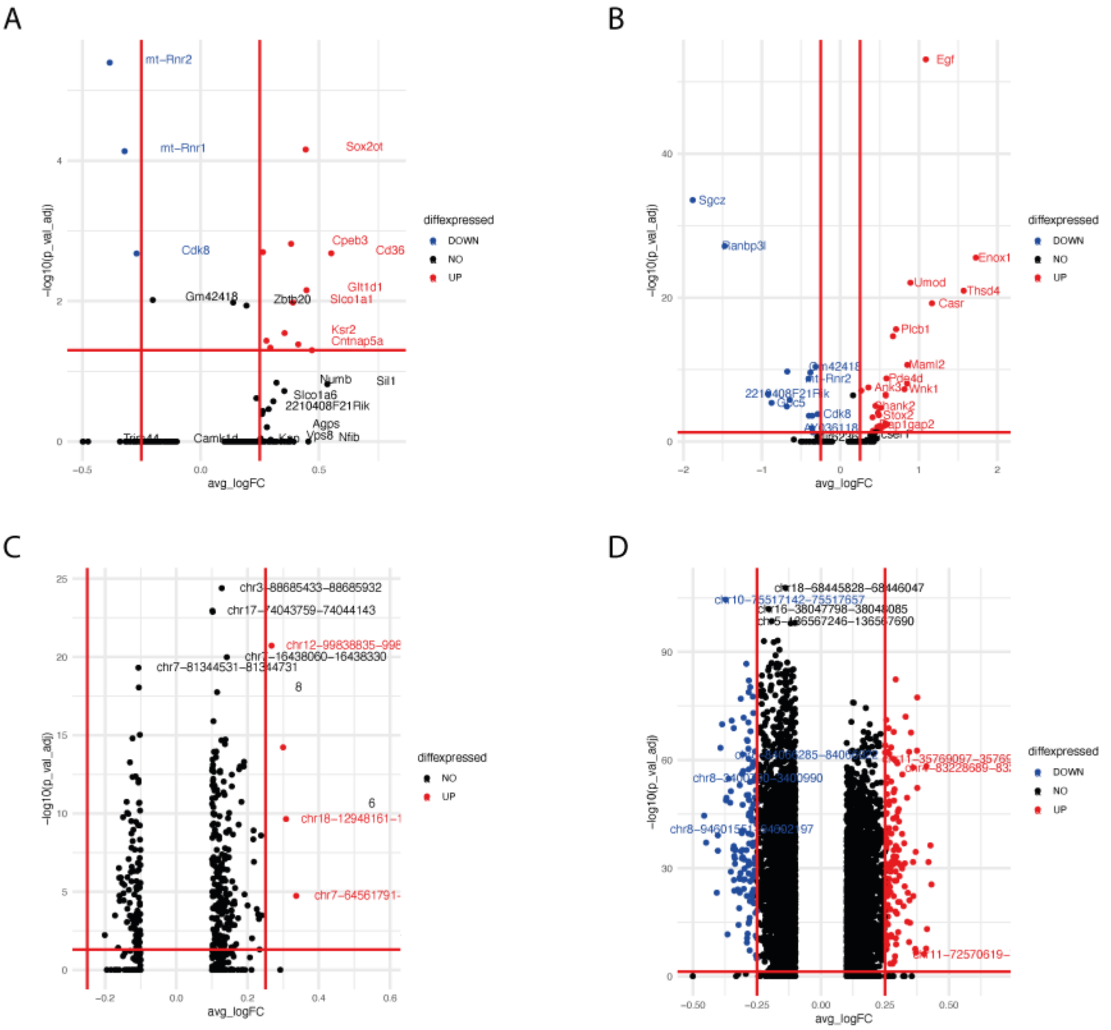


**Supplementary Figure 7 :** Volcano plots showing the differentially expressed genes between (A) cluster 0 vs 13 , and (B) clusters 7 vs 8. (C) and (D) are the volcano plots showing the differentially expressed peaks between cluster 0 vs 13 and 7 vs 8.

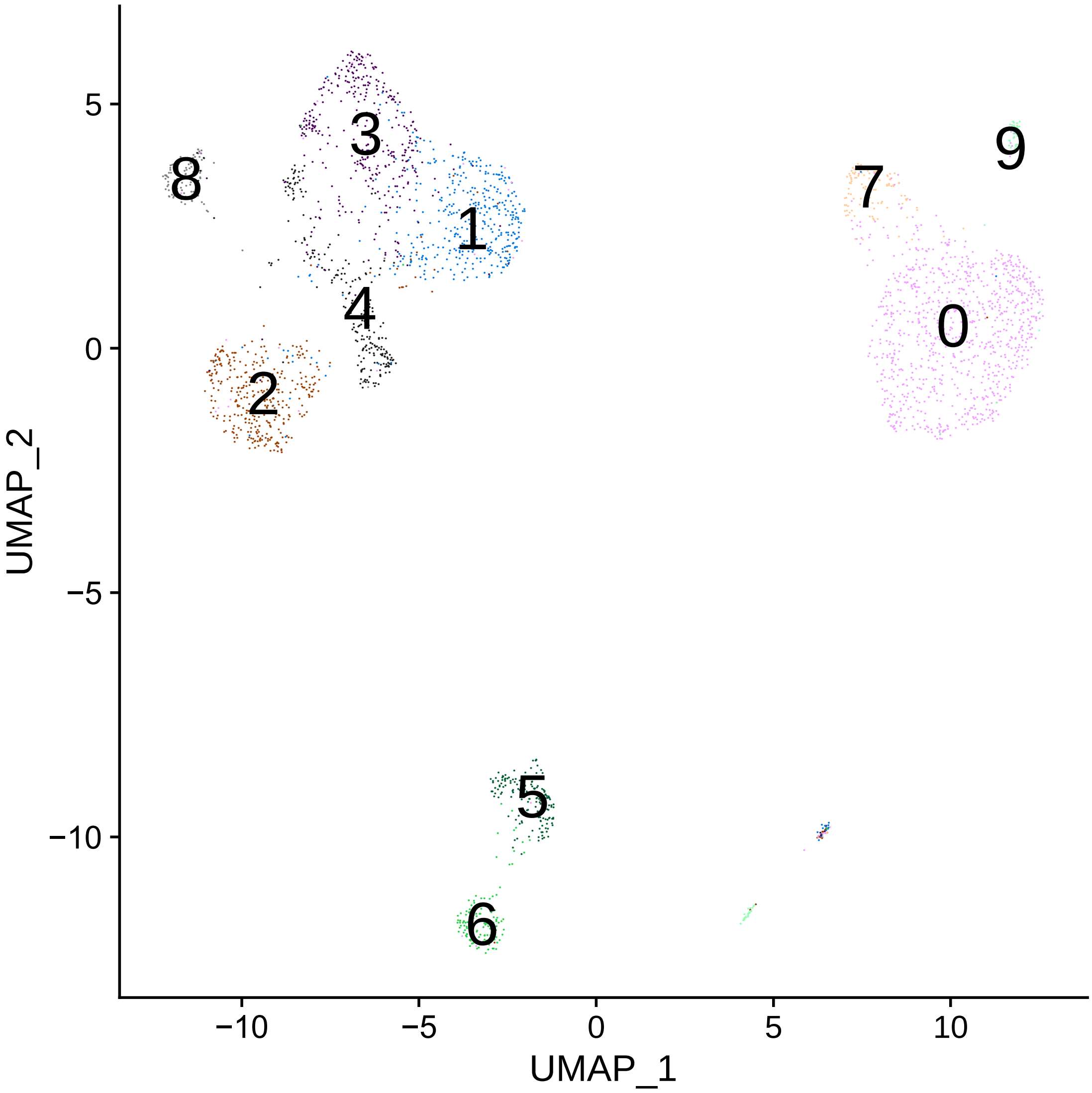


**Supplementary Figure 8:** The 10X genomics multiome data unimputed ATAC cluster overlaid with the RNA clusters. The graph shows the corresponds between the two data domains. In this case the effect of applying scMoC is minimized.

**Supplementary Table 1: Summarizing the limits used in processing different datasets**. *Min genes per cell* defines the almost empty cells, by which cells having less than that threshold of genes is removed. The upper level of genes detected per cell so that the cell is not noisy is set by *Max genes per cell*. *Max Mito percentage* is the maximum number of mitochondrial genes to be accepted in the cell. *Min cells per genes* is set to remove genes that are detected in cells less than this threshold. *Min peaks per cell* and *Max peaks per cell* are the lower and the upper limit of peaks detected in each cell respectively. *Min cells per peak* is set to remove peaks that are detected in cells less than this threshold.

| | sci-CAR | SNARE-seq | 10X genomics |
|---|---|---|---|
| Min genes per cell | 200 | 200 | 100 |
| Max genes per cell | 2500 | 2500 | 5500 |
| Max Mito percentage per cell | 30% | 5% | 30% |
| Min cells per gene | 3 | 3 | 3 |
| Min peaks per cell | 4000 | 4000 | 200 |
| Max peaks per cell | 10000 | 10000 | 32500 |
| Min cells per peak | 3 | 3 | 3 |
